# Habitat remediation followed by managed connectivity reduces unwanted changes in evolutionary trajectory of high extirpation risk populations

**DOI:** 10.1101/2023.11.03.565519

**Authors:** Gina F. Lamka, Janna R. Willoughby

## Abstract

As we continue to convert green spaces into roadways and buildings, connectivity between populations and biodiversity will continue to decline. In threatened and endangered species, this trend is particularly concerning because the cessation of immigration can cause increased inbreeding and loss of genetic diversity, leading to lower adaptability and higher extirpation probabilities in these populations. Unfortunately, monitoring changes in genetic diversity from management actions such as assisted migration and predicting the extent of introduced genetic variation that is needed to prevent extirpation is difficult and costly in situ. Therefore, we designed an agent-based model to link population-wide genetic variability and the influx of unique alleles via immigration to population stability and extirpation outcomes. These models showed that management of connectivity can be critical in restoring at-risk populations and reducing the effects of inbreeding depression; increased connectivity prevented extirpation for the majority of scenarios we considered (71.5% of critically endangered populations and 100% of endangered and vulnerable populations). However, the rescued populations were more similar to the migrant source population (average F_ST_ range 0.05 – 0.10) compared to the historical recipient population (average F_ST_ range 0.23 – 0.37). This means that these management actions not only recovered the populations from the effects of inbreeding depression, but they did so in a way that changed the evolutionary trajectory that was predicted and expected for these populations prior to the population crash. This change was most extreme in populations with the smallest population sizes, which are representative of critically endangered species that could reasonably be considered candidates for restored connectivity or translocation strategies. Understanding how these at-risk populations change in response to varying management interventions has broad implications for the long-term adaptability of these populations and can improve future efforts for protecting locally adapted allele complexes when connectivity is restored.

## INTRODUCTION

Habitat fragmentation is increasing globally, a concern for sustainability because habitat fragmentation impairs ecosystem functions and reduces biodiversity up to 75% (Haddad et al. 2015). As habitat fragmentation continues to encroach on wild environments, connectivity between populations will remain a top priority for population management, particularly for threatened and endangered species due to the often-existing concerns of decreased genetic diversity and increased inbreeding and the potential for these concerns to be exacerbated by increased fragmentation. However, the minimum connectivity required to support population recovery is difficult to predict because the number of migrants needed varies with population characteristics such as extinction risk, migration rate, and genetic makeup of the populations. Here, we quantify these interactions to understand how to maintain genetic variation in recovering populations when repairing corridors or otherwise encouraging movement of genetic variants is desirable.

When genetic diversity is diminished and inbreeding increases, populations often face increased inbreeding depression through the expression of recessive traits and accumulation of maladapted alleles (Coltman and Slate 2003; Del Castillo et al. 2011). Small populations, in particular, are often at high extinction risk due to these effects because they are typically isolated and lose genetic variants quickly through drift (Caughley 1994). Furthermore, populations with high levels of inbreeding depression can enter an extinction vortex when inbreeding depression and genetic drift reduce genetic diversity and fitness in a positive feedback loop (Caughley 1994; Drake et al. 2006; Drake 2008; Braumann 2010; Messer 2016). However, increasing or restoring connectivity can alleviate these stressors by introducing new genetic variants that mask deleterious recessive alleles, thereby decreasing inbreeding depression.

Theoretically, connectivity of subpopulations via one migrant per population per generation is sufficient to retain genetic variation while allowing divergence in allele frequencies; empirically, however, the minimum of one individual per population per generation is often insufficient to achieve neutral or positive population growth for struggling populations (Mills and Allendorf 1996). As such, the genetic connectivity of passing alleles via immigration must also complement demographic connectivity so that the benefits of migrants are reflected in population growth and stability, a goal especially important for management of fragmented and threatened species (Lowe and Allendorf 2010). The benefits of migration – either naturally with corridors or via artificial translocations of individuals from one area to another – depend on the carrying capacity of the source and recipient populations, the rate of migration, the growth rate of the recipient population, the reproductive success of migrants, the competition for resources against other species, and the frequency of repeated migration such that the outcome of immigration events are dependent on examination of these factors before using migration as a conservation tool (Thrimawithana et al. 2013; Backus and Baskett 2021).

Supplementing populations can help sustain populations long term, but at the expense of a decreased effective population size – and therefore decreased diversity in the gene pool – for the first few generations (Ryman-Laikre effect; Ryman and Laikre 1991). This drop in the number of effective breeders in the first generations post-migration arises because the reproductive rate of new individuals tends to be favored, leading to variance in mean fitness between receiving and migrant source lineages (Hagen et al. 2019). This effect is especially pronounced when a greater proportion of the migrating population contains new genetic variants than the receiving population (Christie et al. 2012; Hagen et al. 2020), so identifying the number of new individuals to introduce can be a balancing act between increasing the number of adults while attempting to maintain genetic diversity and fitness. Therefore, establishing if an increase in population size or population fitness is the conservation goal and post-release monitoring of population-wide reproductive success, maintaining an equal sex ratio, and evaluating kinship among offspring is especially important in the first years following assisted migrations.

Because funding is nearly always a limiting factor, most agencies must consider management strategies and the time and resources that population managers must devote to sustaining the species (Waldron et al. 2013; White et al. 2022). Translocation actions for large carnivores can cost at least a few thousand dollars per individual (Weise et al. 2014), with this cost increasing significantly when accounting for unsuccessful reintroductions (Weise et al. 2014). Further, tracking technology, post-translocation monitoring, and other mitigation strategies constitute a higher proportion of project funds than the translocations themselves (Weise et al. 2014; Julien et al 2022) and as a result, some recovery programs choose to focus on data collection rather than action (Buxton et al. 2020). Delaying or shortening the frequency of assisted migration efforts can decrease financial costs, but at a risk of decreased genetic diversity and long-term viability (Martinez-Abrain et al. 2011; Hilbers et al. 2019). Although many have proposed frameworks for species management via translocations (McLachlan et al. 2007; Schwartz and Martin 2013; Williams and Dumroese 2013; Sansilvestri et al. 2015; Barbosa and Tella 2019; Berger-Tal et al. 2019) and for monitoring genetic diversity (Weeks et al. 2011; Furlan et al. 2020; Chen et al. 2021; Hogg et al. 2021; DeWoody et al. 2023), implementation across taxa is still lacking. Thus, identifying how we can most efficiently use funding to support at-risk populations by restoring migration patterns is important for effective management.

Oftentimes, management practices have the goal of both increasing population size and genetic diversity by restoring gene flow via outbreeding with genetically distinct individuals (Weeks et al. 2011; Stepkovitch et al. 2022). Consistently monitoring the genetic diversity of wild populations – both vulnerable and otherwise – before and after management intervention is both costly and difficult *in situ*. Here, we establish a model that can simulate genetic diversity across hundreds of years to evaluate the risks and benefits of increasing migration into high extinction risk populations. Using simulations, we address two questions relevant to understanding evolutionary trends in populations of threatened and endangered species. First we answer the question: How do genetic characteristics of the migrant source population, specifically the population-specific unique genetic variants that determine individual fitness, influence recipient population stability and evolution? We hypothesized that populations receiving migrants with lower heterozygosity (and were accordingly less fit) would be less stable over time compared to populations receiving more fit migrants that have more genetic variants, since low fitness individuals may have reduced effects on the recipient populations’ gene pool. The second question we address is: How does recipient population size (e.g., carrying capacity) and trend (e.g., increasing, decreasing) alter the magnitude of the population’s evolutionary response to immigration? We hypothesized that small recipient populations would represent the migrant gene pools more than moderately sized recipient populations because periods of rapid population growth following an influx of migrants would favor new migrant-brought alleles. These aims will help guide future decisions when considering migration (via corridors or translocations) for managing species on the brink of extinction.

## METHODS

### Demographic model steps

We developed a forward-time agent-based model in R that tracks individuals and their multi-locus genotypes for 350 years (Figure 1). Parameters were selected and evaluated separately to examine their independent and interaction effects (Table 1). We incorporated the effects of migration by creating two populations: a migrant-receiving population and a migrant source population, from which migrants were randomly selected to disperse into the recipient population. In all iterations, the recipient populations were allowed to stabilize for 100 years with a carrying capacity of 1000 individuals (K = 1000) and we then subjected some recipient populations to a 50-year population size reduction to simulate the effects of environmental change; this reduction was comprised of a population size decline (i.e., decreasing trend) over 10 years and a subsequent 40-year period of maintenance at this population size minima. We varied the degree of population size reduction, with three possible reduction proportions corresponding to IUCN (International Union for Conservation of Nature) classifications: vulnerable species had a 30% decline in 10 years (vulnerable carrying capacity = 700 individuals); endangered had a 70% decline in 10 years (endangered carrying capacity = 300 individuals); and critically endangered populations had a 90% decline in 10 years (critically endangered carrying capacity = 100 individuals; Table 1). After the 50-year reduced population period, we simulated habitat restoration by allowing the populations to grow again (i.e., increasing trend), up to the original carrying capacity (1000 individuals), and tracked the populations’ demographic and evolutionary responses for 200 years. Additionally, we included populations that did not face environmental degradation or population decline to serve as controls. Below, we describe the basic model structure and function, and provide further details about each model function in the Supporting Information.

**Figure 1.**
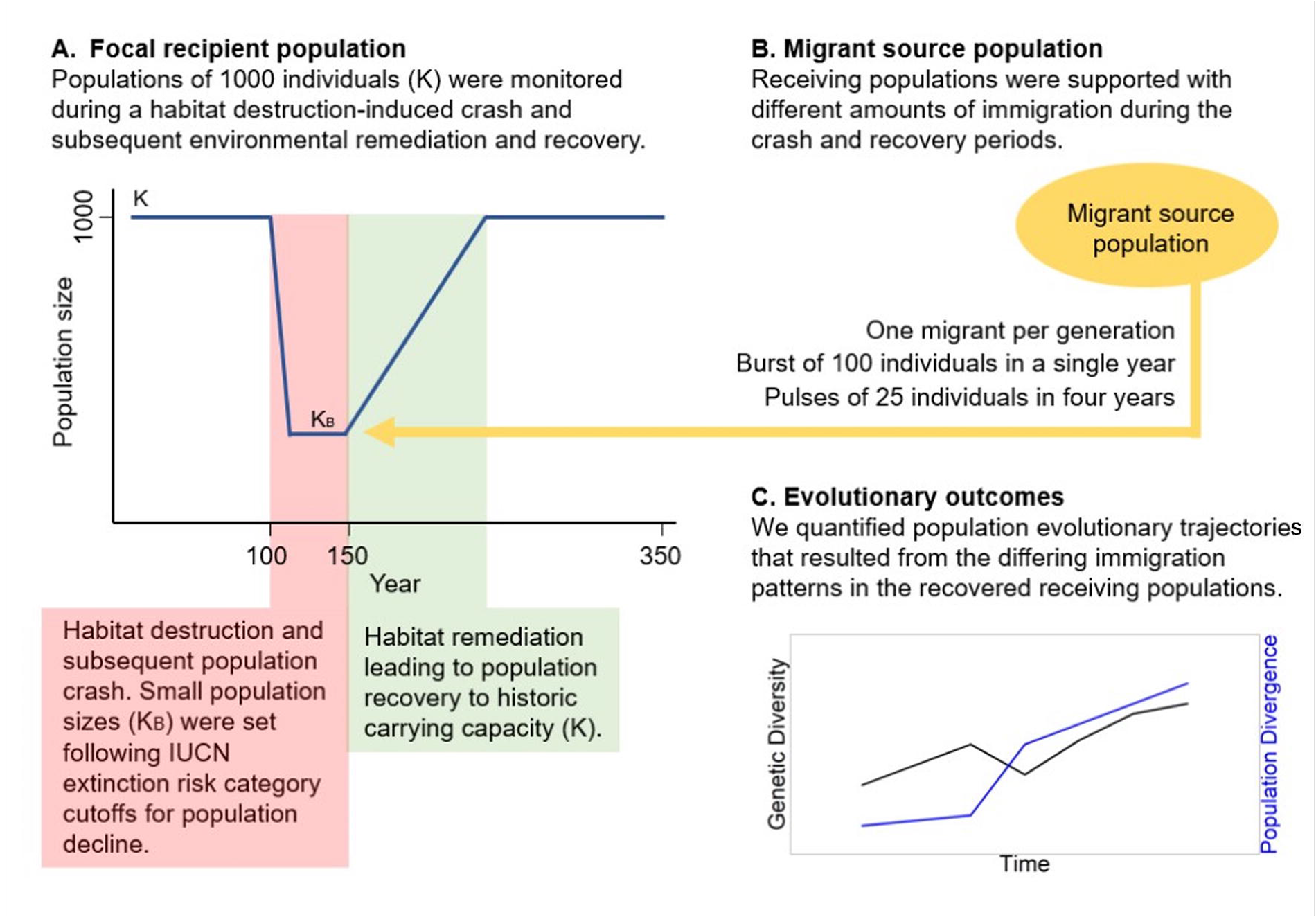
Schematic representation of migrant source and recipient population models. (A) The recipient population was allowed to reach equilibrium and stabilize for 100 years with a carrying capacity of 1000 individuals and was then subjected to a 50-year population decline to simulate environmental change; this population size decline happened over 10 years, and the population was held at the population minima for 40 years at three levels representing IUCN (International Union for Conservation of Nature) and state agency classifications: vulnerable for species that have a > 10% decline in 10 years, K_B_ = 700; endangered with a 50-70% decline in 10 years, K_B_ = 300; and critically endangered with a 80-90% decline in 10 years, K_B_ = 100. After this reduced population period, we simulated habitat restoration by allowing the population to grow again, up to the historical carrying capacity. Additional simulations for populations that did not face population decline were used as a control. (B) Migrants entered the recipient population at a set frequency: 1 migrant per generation, 100 individuals at a single time period (e.g., 100 individuals in year 151) and 25 individuals at four time periods (e.g., 25 individuals in years 151, 165, 181, 195). As a basis for comparison, we also included scenarios where migration was completely absent. (C) We tracked the demographic and evolutionary response of the recipient populations across 350 years.

**Table 1.** Description of parameters and values tested. These parameters describe the starting evolutionary point for the recipient populations, the extent to which population size was reduced to reflect IUCN (International Union for Conservation of Nature) extinction risk designations, and the rate and timing of migration events.

| Parameter | Description | Values tested |
| --- | --- | --- |
| Minor allele frequency | Initial minor allele frequency (p) for migrant source and recipient populations | Low minor allele frequency (0.05-0.15); high minor allele frequency (0.40-0.50) |
| Population crash size | Modified carrying capacity forced on recipient populations from year 100 to 150 to simulate habitat destruction (or degradation); note that census size could be below carrying capacity due to Allee effects and mortality | Carrying capacity (K) reflecting IUCN designations: Vulnerable K = 700, Endangered K = 300, Critically Endangered K = 100 |
| Migration rate | Rate at which individuals moved from migrant source populations into recipient populations | No migration; 1 migrant per generation; single burst of 100 individuals; four pulses of 25 individuals |
| Migrant timing | For burst and pulsed migration, at what point during the population crash and recovery were assisted migrations applied | Migrants were moved to the recipient population during habitat restoration; Migrants were moved to the recipient population only after habitat restoration was completed |

Simulations began by initializing the populations so that the recipient and migrant source populations were at carrying capacity (total recipient population size started at 1000 individuals; total migrant source population size was 5000 individuals) and all individuals were assigned 1100 SNP genotypes. SNPs were split into two types: neutral and migrant-associated. The 1000 neutral SNPs were randomized as either homozygous or heterozygous according to one of two minimum allele frequency ranges for the same allele in both populations and were used for downstream genetic analyses (i.e., 0.05 ≤ p ≤ 0.15 or 0.40 ≤ p ≤ 0.50; Table 1). The 100 migrant-associated SNPs were population specific such that all individuals were homozygous for the source or the alternate recipient population allele and were used to characterize the proportion of migrant ancestry in the population but do not influence fitness. In subsequent generations, each allele had a 0.000001% chance of mutating (µ = 1.0 x 10^-8^), which is similar to empirical estimates of mutation (Nachman and Crowell 2000). Starting individuals were randomly assigned sex and age along a Poisson distribution centered around the age of maturity (1 year). In subsequent years, each surviving individual was aged by 1 year and newly created individuals (see reproduction description below) were assigned genotypes following patterns of Mendelian genetics.

We enacted migration events where random migrants from the source population entered the receiving population and quantified the demographic and evolutionary response of the recipient populations. Within each set of parameter values, migrants entered the recipient populations at a set frequency: 1 migrant per generation, as well as at rates common for assisted migration management (Anderson et al 2005; Kim et al. 2011; Sasmal et al. 2013; Nash et al. 2020; Morris et al. 2021) including 100 individuals at a single time period (i.e., 100 individuals in year 151), which we refer to as burst migration, and 25 individuals at four time periods (i.e., 25 individuals in years 151, 165, 181, 195), which we refer to as pulsed migration. As a basis for comparison, we also included scenarios where migration was completely absent (Table 1). For each assisted migration scenario, we considered the effects of these events both during the population minima when habitat was still being restored (burst immigration year 125; pulse immigration years 125, 140, 155, 170) and after habitat restoration when the population was permitted to grow (burst immigration year 151; pulse immigration years 151, 165, 181, 195; Table 1) to determine if bringing in new individuals before habitat quality is restored would have the same effect on the genetic diversity of the population as bringing new individuals after habitat management and population monitoring occurs.

After the migrants entered the population, individuals were randomly paired for mating and reproduction, and parental pairs that included at least one migrant were preferentially chosen to produce offspring so that the number of migrants closely approximated the number of effective migrants; when migration rates were high and the number of breeding individuals was less than the number of migrants, not all migrants were able to reproduce and, occasionally at small population sizes, migrants did not breed when opposite sex mates were not available. We subsequently considered the effects of this migrant-biased procedure and found that there were similar trends between preferentially choosing migrant parental pairs and completely random mating without a migrant bias (see Supplemental Materials and Supplementary Figure 1). We included population demographic effects in our model to reflect the decreased chance of potential mates interacting with each other at small population sizes (Allee effect; Drake and Kramer 2011) by reducing the chance of finding mates as population sizes decreased; the probability of finding a mate was the complement of the reciprocal of the total number of adult pairs. All mated pairs produced 1-2 offspring, and each year the total number of offspring produced followed the logistic growth equation:

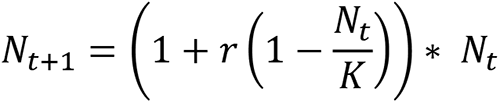

where the population size (*N_t_*_+1_) was determined by the per capita growth rate (*r* = 1), carrying capacity (K = 1000 individuals), and the population size prior to reproduction (*N_t_*). Density independent variation in population size was introduced where the value of *N_t_*_+1_ was randomly generated along a normal distribution with a mean of *N_t_*_+1_ and standard deviation of 1.

Individuals were removed from the population in two ways, one relating to fitness effects and one relating to random environmental or competition effects. To reflect fitness effects, we used the neutral SNPs (i.e., not including source or migrant-receiving population-specific SNPs) to force a fitness cost to reduced genetic diversity; when an individual reached maturity, we removed individuals with a probability equal to the reciprocal of its genomic heterozygosity, such that there was a higher chance of mortality with decreasing heterozygosity. To reflect other causes of mortality in our population, each year we assumed the cumulative probability of death of an individual was equal to the quotient of an individual’s age and the maximum lifespan.

After age nine, individuals were removed from the population through this method. Unless the recipient population size was reduced to 10 or fewer individuals – which we considered functionally extirpated – these events were repeated for 350 years.

### Genetic model steps

We devised several monitoring measures for our recipient populations that persisted across all 350 years. Using the 1000 neutral SNPs, we calculated yearly observed and expected heterozygosity as measures of individual and population-wide genetic diversity by dividing the number of heterozygous neutral alleles over the total number of neutral alleles. We used the migrant-associated SNPs to quantify the proportion of migrant loci over the total number of population-source-associated loci to calculate migrant ancestry in the recipient populations and quantified the number of mating individuals to monitor population fitness. We quantified inbreeding (F_IS_; Supplemental Figure 3) using the *hierfstat* package to examine how immigration of migrants into the recipient populations and random genetic drift changed the progression of the populations. We analyzed divergence (F_ST_) in two ways: the divergence between the historical recipient populations in year 0 compared to each consecutive year of the contemporary recipient populations as well as the historical migrant source populations compared to the contemporary recipient populations in each year using the *hierfstat* package in R (Goudet 2005; Weir and Goudet 2017). Along with the number of effective migrants and the number of effective parents, we also recorded the total number of individuals, sex ratio (Supplemental Figure 4), and number of adults present in each year. The number of migrants in the population and the proportion of migrant-specific alleles in the population served as measures of the demographic and genetic effects of migration on the recipient populations. Finally, the lifetime reproductive success of all individuals was calculated by determining the number of mates, number of offspring, and the number of offspring that survived to maturity for all individuals (Supplemental Figure 5). Significant differences among variables within each year were determined by comparing confidence intervals (84% quantiles approximating α = 0.05; Payton et al. 2003). All scripts and figures were created using RStudio (R version 4.2.1) and all scripts are available at https://github.com/ginalamka/ComplexModel_ABM/.

## RESULTS

Across simulation runs, we found a number of trends that suggest our model was operating within expectations of population genetic theory. For example, during the habitat destruction period, census population size declined to at or below the level as described in IUCN extinction risk designations due to the combination of reduced carrying capacity, Allee effects, and age-related removals (i.e., when K = 1000, N_C_ ≈ 750; Figure 2A). Further, when there was no immigration in our models, increasing the severity of population decline corresponded to proportional decreases in genetic diversity and increases in genetic divergence between historic and contemporary populations (Figure 2). For example, populations with the largest declines in population size (90% decline, critically endangered) resulted in a 52.0% reduction in heterozygosity across neutral SNPs whereas populations with the smallest tested decline (30% decline, vulnerable) had a 14.8% decrease in heterozygosity between years 100 to 350 (Figure 2B). Despite these losses in genetic diversity and the resulting reduced fitness for some individuals, populations returned to pre-bottleneck sizes following habitat restoration (Figure 2A). However, after recovery, populations had diverged substantially from the historical recipient populations and at rates correlated with population crash intensity when the population was small (Figure 1C). These populations also diverged from the historical migrant source population at rates correlating to population crash intensity when the recipient population size was small (Figure 2D). Because there was no migration in this set of analyses, these outputs depict the rate of genetic drift between these two populations. Combined, these outputs illustrate the development of unique evolutionary trajectories that can be altered by population size reductions that ultimately influence individual fitness.

**Figure 2.**
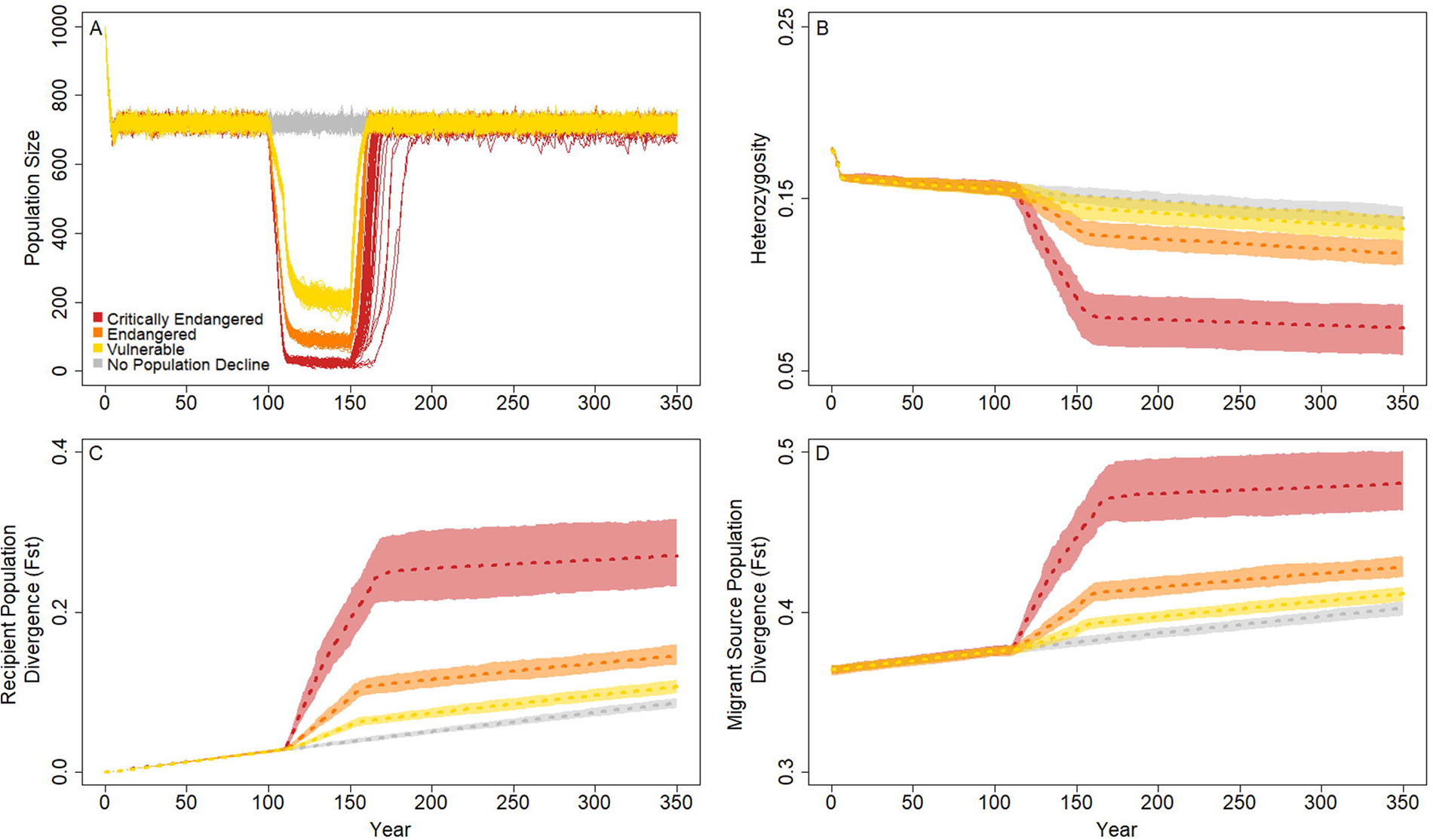
Population genetic and demographic responses in populations compared among extinction risk categories in the absence of migration. The census population size (A) is illustrated for 100 replicates of the simulation runs, where each line represents a single iteration. Here, the census size is depicted as less than carrying capacity to match the point in the model when population response parameters were quantified (i.e., after Allee effects, reproduction, and subsequent mortality). Population response is summarized across these replicates as observed heterozygosity (B), divergence from the historical recipient populations over time (C), and divergence between the recipient and historical migrant source populations at each year (D). In the absence of migration, a drop in census size can push populations on a new evolutionary trajectory with different allele frequencies as a result of the loss of genetic variants that occurred due to drift. Lines in B-D represent mean values across the 100 replicate runs and polygons represent confidence intervals scaled to compare evolutionary outcomes between each parameter set, assuming alpha = 0.05 (i.e., 95% confidence intervals).

### Migrant genetic variance and population stability

We were interested in how migrant-specific genetic characteristics influenced evolution and stability in the migrant-recipient populations. When comparing genetic diversity in populations with a single migrant per generation, we found that starting minor allele frequency (and therefore lower heterozygosity compared to higher initial minor allele frequencies) in the migrant source and recipient populations were not associated with any noticeable effect on average migration ancestry represented in the recipient populations over time (Figure 3A). However, heterozygosity changes in the neutral SNPs differed when starting minor allele frequencies differed; when the source and recipient populations had high heterozygosity initially and no migration, heterozygosity declined by 32.8% over the full 350-year period, although one migrant per generation was effective in slowing this process to a 11.3% loss over the same time period (Figure 3B). Similarly, when the starting minor allele frequencies in both migrant source and recipient populations were small, heterozygosity decreased by 33.6% without migration and was slowed to a 1% loss with migration across the 350 years. When the starting minor allele frequencies in the recipient populations were low but high in the migrant source populations, one migrant per generation increased heterozygosity, particularly through the population decline (years 50-150 resulted in a 70.7% increase due to the influx of new alleles), and ended year 350 with heterozygosities nearest to the populations that started with high minor allele frequencies in both populations with one migrant per generation (H_high > low_ = 0.42 with a 134.7% increase years 1-350, H_high > high_ = 0.44 with a 11.3% decrease years 1-350; Figure 3B). Finally, when the migrant source populations had a low starting allele frequency that migrated into the recipient populations with high initial heterozygosity, the resulting heterozygosity was less than if there were no migrants; migrants with low fitness reduced the recipient population’s heterozygosity by 46.9% (Figure 3B).

**Figure 3.**
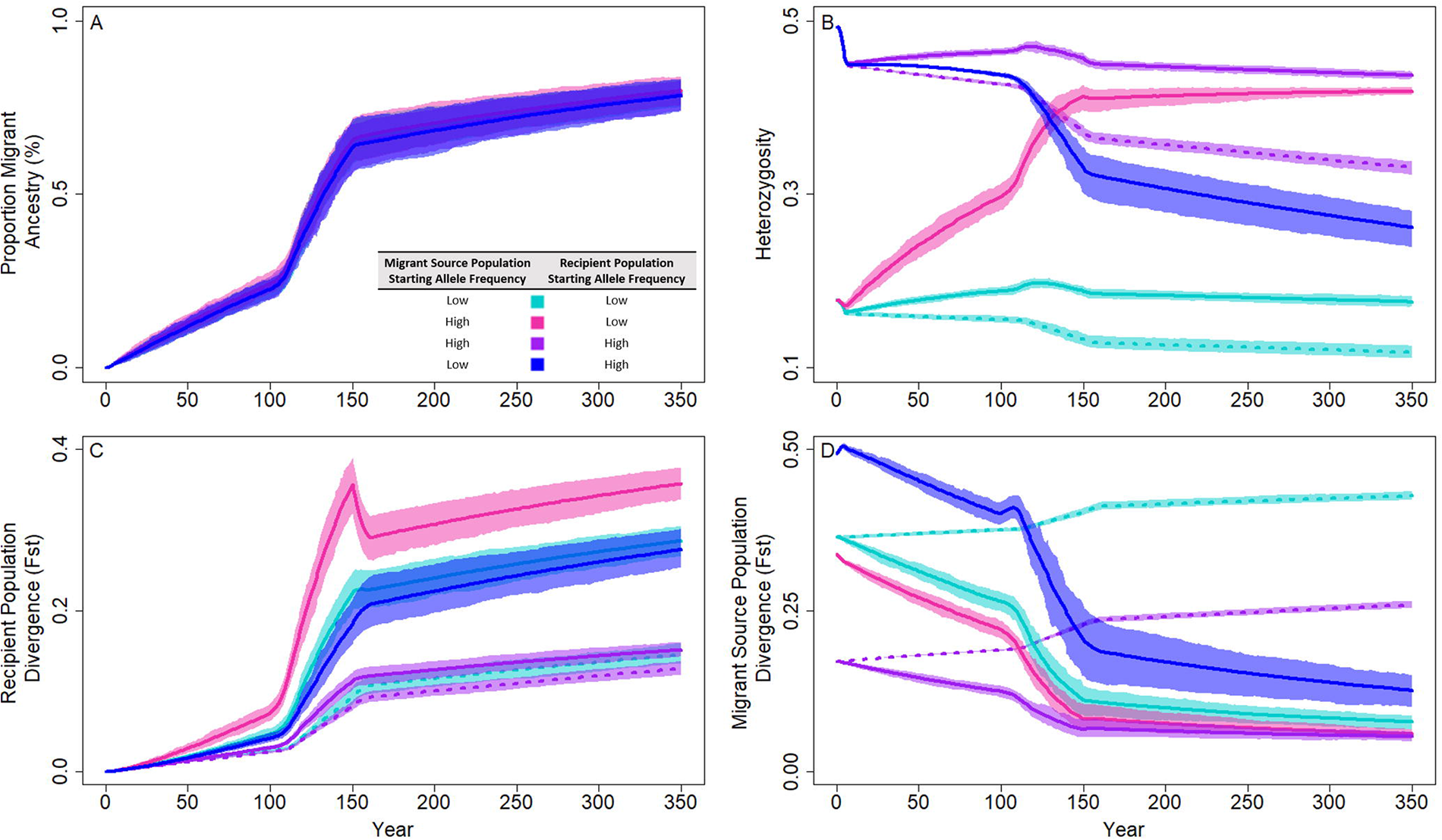
Genetic and demographic responses to migration in populations with different starting minor allele frequencies and a population crash to 70% of the historical population size. Solid lines indicate a single migrant per generation and dotted lines depict scenarios without migration. Starting minor allele frequencies were either low (0.05 – 0.15) in both the migrant source and recipient populations (cyan), high (0.4 – 0.5) in both populations (purple), high in the migrant source populations and low in the receiving populations (pink), or low in the migrant source populations and high in the receiving populations (blue). Lines represent mean values across 100 replicate runs and polygons represent confidence intervals scaled to compare evolutionary outcomes between each parameter set, assuming alpha = 0.05 (i.e., 95% confidence intervals). The proportion of migrant ancestry (A), observed heterozygosity (B), divergence from the historical recipient populations over time (C), and divergence each year from the historical migrant source populations (D) are depicted in each panel. These outcomes suggest that the minor allele frequencies influence evolutionary trajectory of populations connected by migration when migrations occur during a population crash, such that these recipient populations with migrants from populations with higher minor allele frequencies are less diverged from the migrant source populations compared to recipient populations with migrants with lower minor allele frequencies.

We measured divergence (F_ST_) in two ways: the contemporary recipient populations at each year compared to the historical recipient populations and each year between the contemporary recipient and historical source populations. We found that the contemporary recipient populations diverged from the historical recipient populations most slowly when both recipient and source populations had high starting minor allele frequencies and when there was no migration (Figure 3C). When there was a slow influx of migrants and the recipient populations had limited genetic diversity compared to the source populations, the genetic variation of the migrants influenced recipient population divergence, such that the populations diverged significantly faster than all other migrant-recipient population pairs before, during, and after the population size reduction period (depicted as non-overlapping CIs; Figure 3C). This reflects the influx of many new alleles from the migrants into the recipient populations that substantially increased migrant offspring fitness in the contemporary recipient populations. However, all populations with migrants resulted in lower divergence from the historical source populations than in populations without migration; when migrants had lower starting allele frequencies than the recipient populations, the rate of divergence from the historical source populations decreased the fastest, but F_ST_ was still higher than if both populations contained similar starting allele frequencies or when migrants were more fit than the recipient populations (Figure 3D).

### Recipient population size and migration effects

We compared the genetic diversity effects on populations resulting from a single migrant per generation to no migration when the migrant recipient populations were reduced to one of three extinction risk levels. Despite a brief period of decline in genetic diversity when the population size dropped (years 100 – 150), we found that heterozygosity was maintained in all combinations of migration and population crash sizes (years 150 – 350; Figure 4B). Comparing within each extinction risk category, we found that connected populations tended to be more genetically diverse than the unconnected populations. However, even when connected by one migrant per generation, populations reduced to the critically endangered size had similar heterozygosity as unconnected vulnerable and unconnected non-bottlenecked populations (evidenced by overlapping CIs; Figure 4B). These trends were driven directly by the increasing influence of the migrants; over time, the proportion of migrant ancestry in the recipient populations increased in all populations regardless of the magnitude of the population crash (i.e., the relative extinction risk categories), but populations with the largest declines ended with the largest proportion of migrant ancestry (nearly 100% in the critically endangered populations; Figure 4A). Because the single-migrant populations with lower extinction risk lost fewer alleles during the population crash, these populations were less penetrable by new migrant-brought alleles, meaning that migrant ancestry accumulated less in these populations.

**Figure 4.**
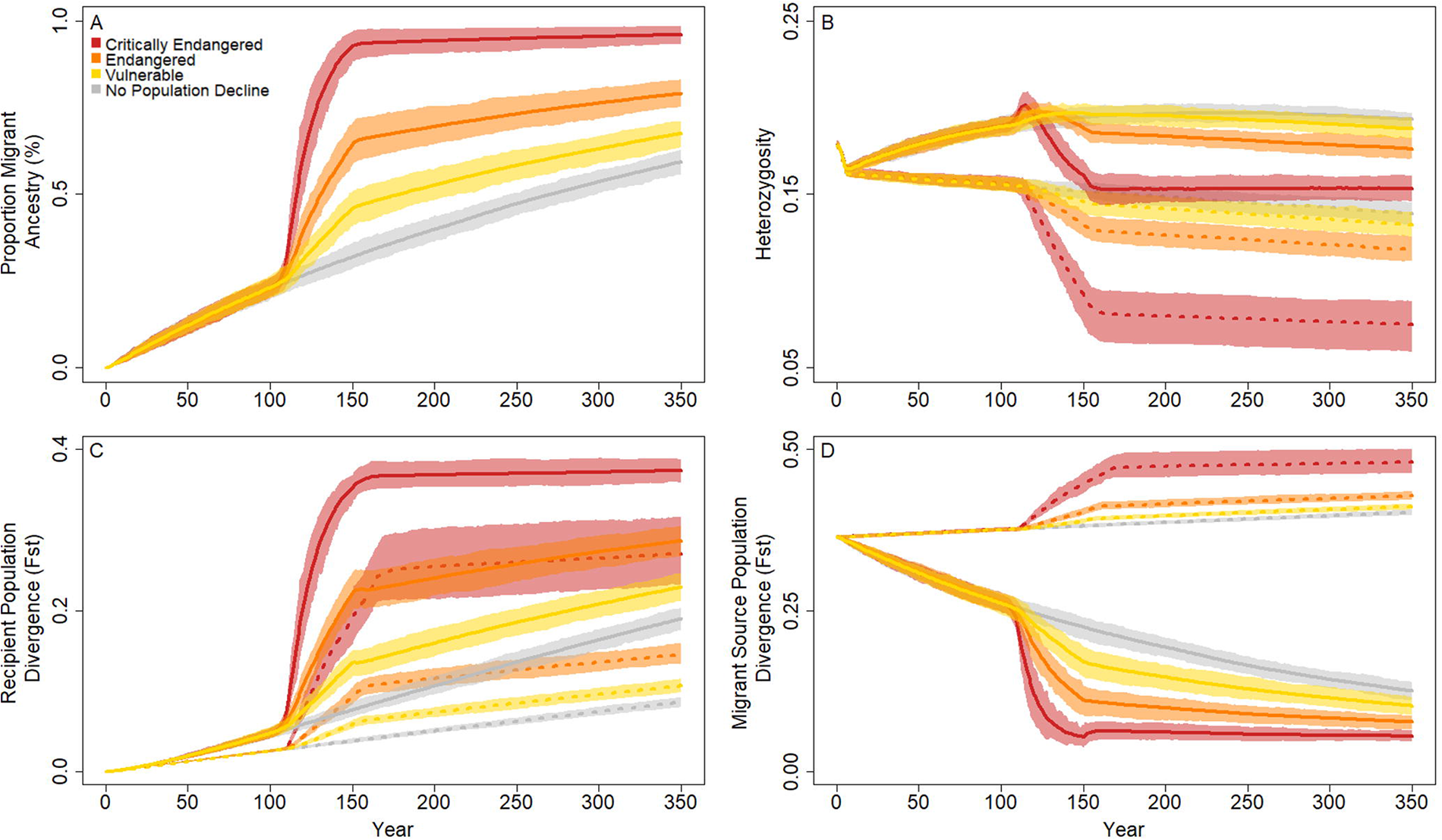
Population genetic and demographic outcomes of incorporating migration into population crash recovery. Populations were shrunk (years 100-150) to sizes reflecting common extinction risk classifications (critically endangered, 90% reduction; endangered, 70% reduction; vulnerable, 30% reduction; no population reduction), and movement of a single individual per year (solid line) was used to support and bolster these populations through remediation (year 151) and population recovery. These trends were compared to the same demographic patterns but without migration (dotted lines). The proportion of migrant ancestry present in the recipient populations (A), observed heterozygosity (B), divergence of the recipient populations from the historical populations over time (C), and divergence of the recipient populations from the migrant source populations each year (D) illustrate the new evolutionary trends resulting from these migration decisions. Lines represent mean values across 100 replicates and shaded areas represent the confidence intervals needed to compare among parameter sets assuming alpha = 0.05 (i.e., 95% confidence intervals).

These patterns of differences among extinction risk categories related to migrant population ancestry was also reflected in population divergence (F_ST_) trends, where unconnected populations diverged less from the historical recipient populations than their connected recipient population counterparts (Figure 4C). In addition, the severity of the population crash was inversely proportional to the strength of divergence observed; at the end of the 350-year observation period, mean F_ST_ between the historical recipient populations and the contemporary recipient populations at year 350 ranged from 0.23 in the vulnerable populations to 0.37 in the critically endangered populations when connected by one migrant per generation and from 0.11 in the vulnerable populations to 0.27 in the critically endangered populations in the absence of migration (Figure 4C). When comparing divergence of the recipient populations to the historical migrant source populations, we found that connected and unconnected populations had differing trends; unconnected populations drifted from the historical migrant source populations whereas the connected recipient populations tended to become more similar, with the size of the population crash relating to the magnitude of this divergence. At the end of the year 350, mean F_ST_ between the historical migrant source populations and the contemporary connected recipient populations ranged from 0.05 to 0.10 and the unconnected populations ranged from 0.41 to 0.48 in the critically endangered and vulnerable populations, respectively (Figure 4D). Although the range between the minimum and maximum F_ST_ within the connected and unconnected populations was similar, the connected populations moved much further from the initial starting value (F_ST_ = 0.36) compared to the unconnected populations, indicating accelerated divergence in the populations connected by migration compared to the populations without migration.

We evaluated changes in genetic diversity, accumulation of migrant ancestry, and population divergence among the four migration scenarios (one migrant per generation, burst of 100 migrants at a single time point, pulse of 25 migrants each at four time points, no migrants) within each extinction risk category and in the absence of a population crash (Figure 5). Trends among migration scenarios within each extinction risk category were similar, with the largest within-group differences in the critically endangered category (90% decline); heterozygosity remained consistent over each time step except during the population crash (years 100 to 150; Figure 5D-5F). Within each extinction risk category, burst and pulsed migration scenarios resulted in similar heterozygosities, surpassing heterozygosity in the one migrant per generation scenario only in critically endangered populations as the population size increased, with final heterozygosity similar in all populations with migrants in all extinction risk groups by year 350 (i.e., overlapping confidence intervals). In contrast to populations that contained migrants, the final mean heterozygosity was significantly lower in the no migration scenarios in all extinction risk groups (Figure 5D-5F). These trends were driven by migrant contributions to these populations; as the proportion of migrant SNPs represented in the recipient populations increased, neutral alleles originating in the migrant source populations also increased in frequency. Populations connected by a single migrant per generation had a larger proportion of alleles with migrant ancestry in most generations compared among all other migration schemes and within each population crash size category (Figure 5A-5C). As individuals migrated and the contemporary recipient populations increased to its historic size, alleles with migrant ancestry were maintained due to the increased fitness of these lineages relative to some of the existing families in each population (i.e., increased heterozygosity due to increased fitness in migrant-resident pair offspring compared to offspring with both resident parents). Furthermore, the burst migration scheme resulted in increased migrant ancestry compared to the pulsed scheme; in the endangered population crash size category, for example, the burst introduction of 100 individuals resulted in 10% more migrant ancestry than pulsed migration (38% vs 28% migrant ancestry years 200 – 350). Thus, even though heterozygosity was equivalent in burst and pulse populations from years 200 – 350, the genetic variants had different sources suggesting the potential for different evolutionary and ecological outcomes.

**Figure 5.**
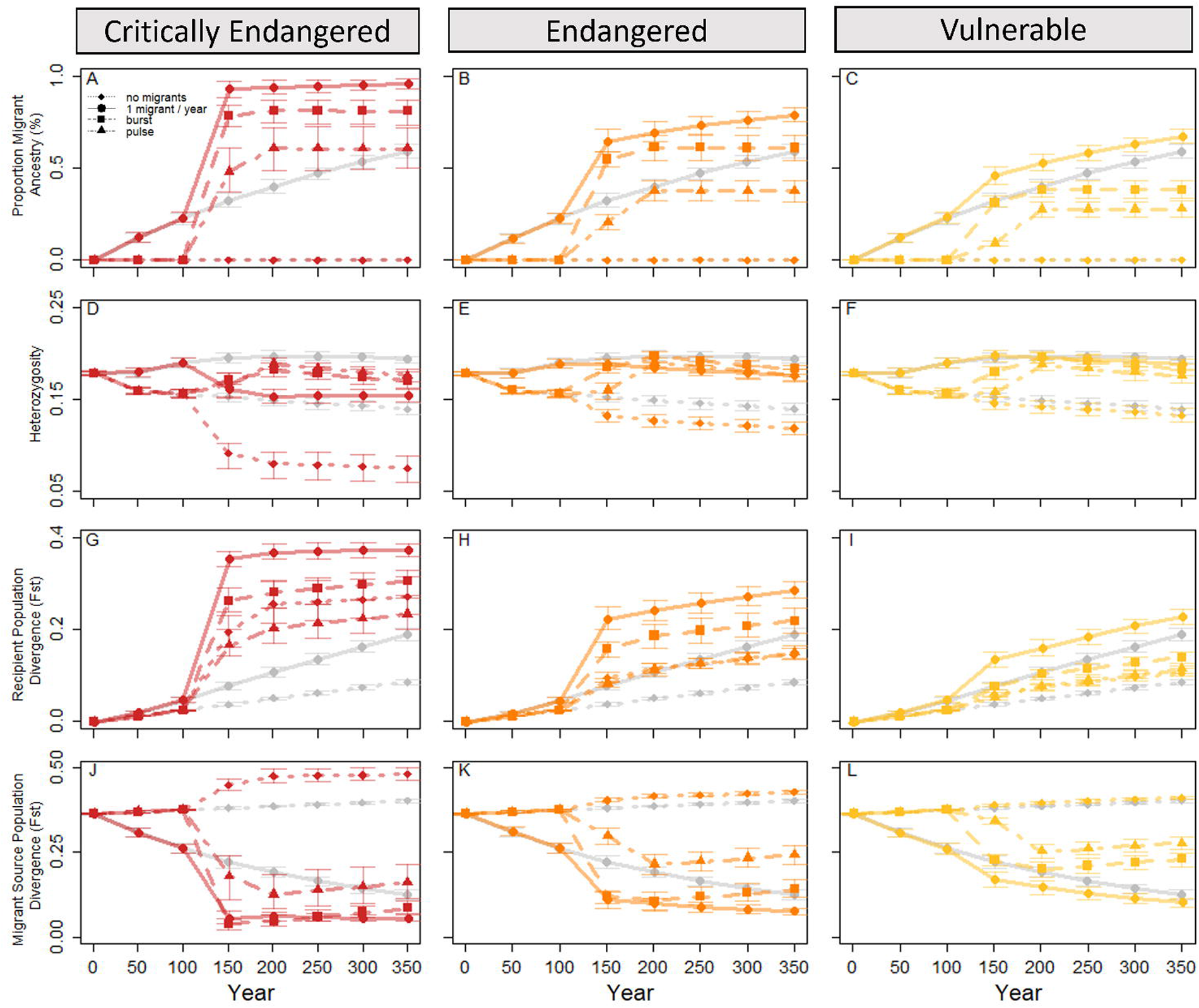
Population genetic and demographic outcomes in populations compared among extinction risk categories while incorporating migration into population crash recovery. Populations were shrunk (years 100-150) to sizes reflecting common extinction risk classifications (critically endangered, 90% reduction; endangered, 70% reduction; vulnerable 30% reduction; no population reduction), and movement of a single individual per year (solid line), burst migration of 100 individuals once (year 151; dashed line), and four pulse migrations of 25 individuals (years 151, 165, 181, 195; dashed and dotted line) was used to support and bolster these populations through remediation (year 151) and population recovery. These trends were compared to the same demographic patterns but without migration (dotted line) and in the absence of a population crash (grey lines). The proportion of migrant ancestry present in the recipient populations (A-C), observed heterozygosity (D-F), divergence of the recipient populations from the historical population over time (G-I), and divergence of the recipient populations from the historical migrant source populations each year (J-L) illustrate the new evolutionary trends resulting from these management decisions every 50 years. Lines represent mean values across 100 replicates and error bars represent the confidence intervals needed to compare among parameter sets assuming alpha = 0.05 (i.e., 95% confidence intervals).

Overall, divergence increased between the contemporary recipient populations and the historical recipient populations over time (Figure 5G-5I) and decreased between the contemporary recipient populations and historical migrant source populations each year (except in the absence of migration; Figure 5J-5L). As populations returned to their historic sizes, pulsed migrations diverged from the initial population the least, followed by no migration, burst, and single-individual migration diverged the most, with magnitudes for each related to the size of the population crash (i.e., larger percent population reduction resulted in more divergence). These divergence patterns contrast with divergence of the recipient populations to the historical migrant source populations; for all population crash categories, the populations not connected by migration drifted away from the historical migrant source populations. All three sets of migrant connected populations ended more similar to the historical migrant source populations than they started, with a single migrant per generation and burst migrations being the most similar to the migrant source populations immediately after habitat recovery (year 150) and pulsed migration being significantly less similar that same year. After assisted migrations concluded, the burst and pulsed migration populations diverged away from the historical migrant source populations along their own evolutionary trajectories. Comparing across extinction risk categories and within migration schemes, the smaller the population minima, the more diverged these populations were from the historical recipient populations (Figure 5G-I) and the more similar to the historical migrant source populations (Figure 5J-5L).

Consistent with trends comparing unconnected populations to populations connected by a single migrant per generation, burst and pulsed migrations resulted in population genetic and divergence outcomes that scaled with the severity of the population crash. For example, in both burst and pulsed migrations, critically endangered populations resulted in a higher proportion of migrant SNPS and were more diverged from the historical recipient populations than other extinction risk categories (Figure 5). Similarly, these populations were less diverged from the historical migrant source populations than the other population categories under the burst and pulse migration model. Combined, these trends represent the effects of migrant ancestry that build up at rates proportional to the recipient population size as long as migration continues.

Finally, we considered how the timing of assisted migration events affected genetic diversity of the recipient population. Specifically, we compared population outcomes when migration occurred during population minima (and therefore during habitat restoration) against when migration occurred during recovery phases (after habitat restoration; Figure 6). Burst migrations resulted in similar outcomes between these two timing scenarios, although heterozygosity was slightly higher and divergence from the historical recipient populations slightly lower when migration was initiated during the recovery phase (Figure 6A and 6E). In contrast to this, pulse migration that occurred prior to the population recovery period resulted in almost double the frequency of migrant alleles compared to if migration had occurred after population recovery (67% vs. 38%; Figure 6B). As a result, pulsed migration that happens during population recovery resulted in populations that more closely resembled the historical recipient populations and were more diverged from the historical migrant source populations in year 350 compared to when migration occurred before the population started growing (Figure 6F and 6H).

**Figure 6.**
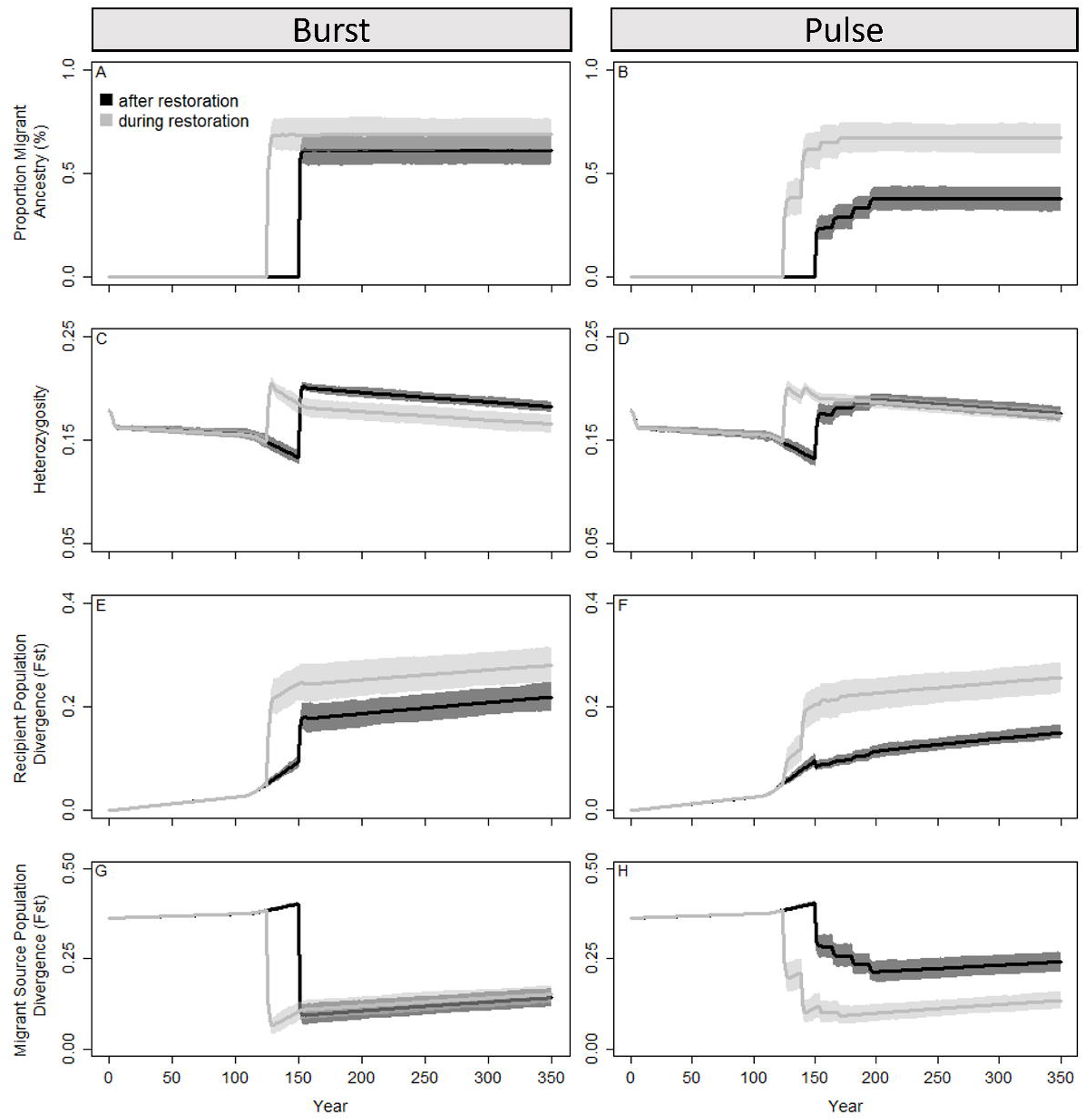
Population genetic and demographic responses with a population crash to 70% of the starting population with different timings of management intervention. Assisted migrations were either burst translocations (100 individuals once) or pulsed (25 individuals 4 times). Light grey lines depict assisted migrations implemented concurrently with habitat restoration (year 125 for burst; years 125, 140, 155, and 170 for pulse) and black lines indicate migrations that occurred after habitat restoration was completed (year 151 for burst; years 151, 165, 181, and 195 for pulse). Lines represent mean values across 100 replicate runs and polygons represent confidence intervals scaled to compare evolutionary outcomes between each parameter set, assuming alpha = 0.05 (i.e., 95% confidence intervals). The proportion of migrant ancestry (A), observed heterozygosity (B), divergence from the historical recipient populations over time (C), and divergence each year from the historical migrant source populations (D) are depicted in each panel. These outcomes suggest that implementing a pulse migration after restoration and with population growth will supplement populations with similar levels of increased genetic diversity and with less influence of alleles with migrant ancestry.

## DISCUSSION

Although one migrant per population per generation is regarded often as the ideal population connectivity scenario because higher genetic diversity is generally predictive of less inbreeding and more stable populations (DeWoody et al. 2021; Kardos et al. 2021), connectivity can have unintended effects if local alleles that are swamped by migrant variants ultimately reduce fitness (Lenormand 2002). Here, we consider the contrasting effects of inbreeding depression and connectivity in the context of conservation by modeling the effects of migration on recovery of at-risk populations. Specifically, we consider populations from critically endangered, endangered, and vulnerable species, vary migration rates and frequencies as well as migrant source population characteristics, and consider the long-term effects that various management decisions can have on these conservation targets. Overall, we find that the timing and rate of migration is a critical predictor of the genetic make-up of recovered populations and that some combinations of these parameters can send newly connected populations on entirely new evolutionary trajectories. Through this work, we add to the body of evidence that identifies habitat suitability, management actions, and population connectivity as the most limiting factors for a species’ long-term viability (Bouzat et al. 2009; Baling et al. 2016; Bubac et al. 2019; Hilty et al. 2020; Goicolea et al. 2022; Gonzalez et al. 2023; Li et al. 2023; Smith et al. 2023) and provide a model that can evaluate the risks and benefits of common management strategies.

Similar to other theoretical and empirical research relating the severity of population crashes to population viability and the ability of migration and selection to effectively swamp local alleles (Fagan and Holmes 2005; Lenormand 2002), here we show mechanistically how extinction risk (i.e., population size) influences the relationship between these forces; our model shows that reduced heterozygosity, high divergence from local alleles, and high relatedness to migrant source populations occurs in connected populations with high intensity crashes. Ultimately, predicting the populations’ response to a single migrant per generation or assisted burst or pulsed migrations depends on the system and goal of management interventions. For example, gene flow can constrain adaptation to local conditions by passing on migrant derived traits, potentially lowering fitness via outbreeding depression when fecundity is low or turnover is high (Frankham et al. 2011; Weeks et al. 2011). In our model, accumulation of migrant ancestry alleles occurred at a faster rate with low frequency but constant immigration as compared to burst or pulsed assisted migration strategies, sometimes resulting in lower genetic diversity in the long term (Figure 5). However, migrations also act as direct support in preventing extirpation that, when it occurs, removes many of the locally adapted allele complexes. Thus, balancing the risks of increased connectivity with the risks of complete population loss is tied to population-specific pressures and management goals.

We investigated the effects of the average minor allele frequency in the migrant source and recipient populations as a potential avenue for lessening the migration risks to populations on the edge of extirpation (Figure 3). This is based on the idea that migration and selection can be balanced across populations and gene variants will be maintained in the recipient populations even with differing population-wide fitness (Lenormand 2002). In lab-based mesocosms, Trinidadian guppy populations with lower starting genetic variation and smaller effective population sizes benefitted from gene flow more than populations with higher diversity and larger sizes, suggesting that evolutionary history can be predictive of individual and population level fitness (Miller et al. 2019). Although the accumulation of migrant-derived alleles was similar in the high and low minor allele frequency comparisons, the divergence from the historical recipient populations was greatest when one or both migrant-connected populations had a low starting allele frequency and divergence from the historical migrant source populations was smallest when migrants have high starting allele frequencies. Because populations with high conservation need are often small and have reduced genetic diversity, this supports the idea that identifying populations with similar allelic variants, especially putatively adaptive variants, may provide an avenue for demographic support with less risk to the existing population (Weeks et al. 2011; La Haye et al. 2017; Bertola et al. 2022). Here, we underscore the importance of this consideration as our model suggests that the neutral portions of recipient populations are unlikely to return to their historical genetic composition once migration ends (Figure 3C and 3D).

Maintaining behavioral, morphological, and metabolic adaptations expressed via locally adapted alleles is often an important consideration in introducing gene flow among populations. Although one migrant per generation can result in increased heterozygosity over time, the tradeoff is that approximately 50-100% migrant derived alleles will drive this increase in diversity, resulting in a population that more closely resembles the migrant source population – and therefore retains migrant-favored traits – rather than traits that have accumulated in the historical recipient population (Figure 4). Increased migration and selection in favor of migrant traits further drives these demographic responses, which occurs at faster time scales than genetic drift (see Supplementary Figure 1), despite the effect of this relationship being proportional to population crash sizes. In the case of rare species, maintaining these derived traits is imperative to the sustainability of the species.

The inverse relationship between genetic diversity and the presence of admixed alleles is especially evident in populations that exhibit introgression and hybridization (Allendorf et al. 2001); backcrossing of previously isolated populations homogenizes historically unique alleles. While homogenization can contribute to the loss of rare traits, introgression can support populations in rapidly changing environments that require more than the current standing genetic diversity (Benjamin-Fink and Reilly 2017; Ranke et al. 2020). Unfortunately, until admixture occurs in wild populations, it is hard to predict if hybridization of distinct populations will result in more adaptive phenotypes (potentially due to increased heterozygosity) or disruption of co-adapted gene complexes.

Assisted migration, such as the burst and pulse migrations simulated here, develop alternative evolutionary trajectories, evidenced by a similar increase in heterozygosity but different rates of decreased divergence from the migrant source populations. Burst and pulse migrations result in overall similar heterozygosities across all extinction risk categories, but a slower retention of migrant derived alleles in pulsed migrations, depicting a lessened Ryman-Laikre effect in this management plan (Figure 5). Therefore, pulse migrations seem to be more effective than burst migrations for increasing diversity and population size while maintaining local alleles. For example, in an endangered *Cricetus cricetus* population, genetic diversity initially decreased after the first translocation when reintroduced individuals were genetically different from the receiving population, but additional supplementation increased genetic diversity regardless of the initial diversity of the source and recipient populations (La Haye et al. 2017). This disparate adaptive potential may be due to alternative responses to selection; gene-frequency estimates (e.g., heterozygosity) responds to selection in the short-term while allelic-diversity estimates (e.g., migrant ancestry) reflect long-term adaptation and total selection responses because segregating sites within a subdivided population are more strongly constrained by natural selection than neutral alleles (Cabellero and García-Dorado 2013). In addition to the genetic benefits of pulse migrations, when management utilizes multiple translocations, recruitment of migrants into the population may increase via increased learning of newly translocated individuals from those surviving from a previous reintroduction, higher availability of density-dependent resources, relaxed Allee effects, a buffer against high mortality from stochastic natural disturbance, and feasibility of more frequent but smaller management actions (IUCN/SCC 2013; La Haye et al. 2017; Martin et al. 2017; Lewis et al. 2022).

Just as the frequency of introductions alters the demographic and genetic effects in the receiving population, we examined how the timing of translocations influences the accumulation of traits with migrant ancestry. When migrations occur before the habitat is restored and while the population remains at low population sizes, accumulation of migrant traits and increased divergence from the historical population occurs long-term, altering the demographic and evolutionary trajectories compared to if the habitat quality increased and the population was permitted to grow post restoration (Figure 6). Waiting to begin translocations until habitat quality increases is an effective management strategy (see Figure 2 in Martin et al. 2017), as long as the population is not extirpated during the waiting period. In our system, critically endangered populations without gene flow were eradicated within 18-65 generations of population decline commencement (Supplementary Figure 2), which could be before assisted migrations are implemented if the habitat is unsuitable for supplementation. Though gene flow can temporarily increase population viability, without addressing the cause of the decline — often habitat destruction — the long-term sustainability is still at risk (Grant et al. 2019). While translocations were and still often are regarded as the last-ditch effort due to long-lasting effects on the population and environment, we recommend active and proactive restoration efforts over natural regeneration (Watson and Watson 2015), especially for populations with high risk of extirpation and when proper techniques and genetic considerations are implemented (Chipman et al. 2008; Weeks et al. 2011; Berger-Tal et al. 2019; Morris et al. 2021).

Of course, instead of active population management and monitoring, many of these difficulties in assisted migrations could be alleviated with increased habitat quality and decreased fragmentation; corridors that allow for gene flow among populations are an effective strategy that can boost viability (Sharma et al. 2013; Christie and Knowles 2015; Hevroy et al. 2017; this study). Although unassisted gene flow via habitat restoration and connectivity could be more time and cost effective long-term than repeated assisted gene flow, the desired genetic diversity, relatedness of migrant source and recipient populations, and the desired rate of population growth must be considered. Indeed, our simulations suggest that populations with connectivity via one migrant per generation are slower to grow in size and retain migrant alleles faster than populations with assisted migrations but can still be more useful for conservation planning than continued population fragmentation. Additional investigations into the adaptability of populations, perhaps through epigenomic modifications or selection based on changing environmental conditions could be additionally useful in population management. Further, examination of other genetic problems that are reflected in small populations like mutational load, inbreeding depression, and fixation of maladaptive alleles are promising avenues for future viability analyses. Ultimately, protecting and maintaining species at the brink of extinction will benefit biodiversity and the community by building engagement and collaboration (Kremen and Merenlender 2018; Zellmer and Goto 2022), in addition to the genetic and demographic benefits of connected populations to species sustainability.

Data Archiving Statement: All scripts and data for this study are available at https://github.com/ginalamka/ComplexModel_ABM/.

