## Supplemental Information for "Habitat remediation followed by managed connectivity reduces unwanted changes in evolutionary trajectory of high extirpation risk populations"

**Supplementary Methods**

We created an agent-based model to predict long term evolutionary trends by introducing various migration rates in populations with varying levels of extinction risk. Below, we provide additional information about each model function.

Initialize recipient and source populations

Simulations began with the recipient population at carrying capacity (K = 1000) and a large source population ($K_S$ = 5000). Individuals were assigned an individualized ID number, age according to a Poisson distribution (rpois function in R) minus one, and a random sex (0 = female, 1 = male). Additionally, all individuals were characterized by two sets of SNPs: neutral and assigned with varying minor allele frequencies ($nSNP_D$ = 1000; $0.05 \leq p \leq 0.15$ or $0.40 \leq p \leq 0.50$) and fixed for different alleles in each population ($nSNP_M$ = 100) so that the recipient population was homozygous for one allele and the source population was homozygous for the alternative allele. Therefore, a total of 2200 alleles describe each individual, with each locus defined as heterozygous (01) or homozygous (either 00 or 11). Relative fitness, calculated as the observed heterozygosity across the $nSNP_D$ loci was calculated for all individuals. The population then went through each of the life history events described below yearly for a total of 350 years.

Age Up

The first step in the model involved incrementing the age of all surviving individuals by one year. Because this is the first step in the model, the initialized population consisted of individuals that would be "born" in the first year.

Death

There were two functions that removed individuals from the population. Following the aging of individuals, a fitness-induced chance of death was imposed on the individual at the age of maturity equal to the inverse of the observed heterozygosity of the individual so that individuals with a greater heterozygosity had a greater fitness and a decreased chance of death. Additionally, as the last step in each simulation year, we assumed the cumulative probability of death of an individual was equal to the quotient of an individual's age and the maximum lifespan. Individuals that were forced into mortality were removed from subsequent model steps.

Migrate

Migrants were randomly selected from the source population to move into the recipient population, with the number and timing of migrants per generation selected determined based on the parameter set. Although migrants move into the recipient population and were preferentially chosen for mating pairs, the number of migrants in the population are not exactly equal to the number of effective migrants in that generation since the number of migrant pairs could be more than the number of offspring generated in that generation or, at small population sizes, opposite sex individuals may not be available.

Mate Choice

Reproduction occurred between randomly selected pairs of adults, with preference given to migrants. Parents of the opposite sex were matched as mates with replacement so that individuals could mate with more than one individual in that year. An Allee effect was imposed so that as the number of individuals in the population decreased, the chance of mates interacting decreased; the probability of finding a mate was the complement of the reciprocal of the total number of adult pairs.

Setting Population Size

The number of offspring produced per year was determined using the logistic growth equation so that the total number of individuals in the next year was calculated. The recipient population was allowed to persist around carrying capacity for the first 100 years. Following that 100-year period, the population was forced through a bottleneck for 10 years, persisted at a lower carrying capacity for 40 years, and then allowed to expand at a rate of logistic growth up to the original carrying capacity of 1000 individuals. Logistic growth was calculated with the equation

$$N_{t+1} = \left(1 + r\left(1 - \frac{N_t}{K}\right)\right) * N_t \qquad (1)$$

where the population size ($N_{t+1}$) was determined by the per capita growth rate ($r$ = 1), carrying capacity (K = 1000), and the population size prior to reproduction ($N_t$). The size of the carrying capacity at the duration of the bottleneck ($K_B$) was quantified so that the new carrying capacity would be within limits for IUCN criteria for the evaluation for the Red List. Specifically, IUCN criteria evaluate species that are vulnerable as those that have > 10% decline in 10 years ($K_B$ = 700), endangered with a 50-70% decline in 10 years ($K_B$ = 300), and critically endangered with a 80-90% decline in 10 years ($K_B$ = 100).

Reproduction

Offspring genotypes were assigned according to Mendelian inheritance, where one allele at each locus was randomly selected from each parent. Mutation on generated genotypes was at a rate of $1.0 \times 10^{-8}$ mutant/generation/allele. If an allele was selected to mutate, the allele was switched from either a 0 to a 1 or a 1 to a 0, depending on the initial identity.

Lifetime Reproductive Success

Prior to migration of individuals into the recipient population, we assumed there was no prior history of reproductive success for migrants. Therefore, at the conclusion of each replicate of the simulation, lifetime reproductive success was calculated by determining the sum of offspring that were generated and survived to maturity by each parent while living in the recipient population.

Analyze

After each generation, we calculated population demographics to monitor the population. Demographic calculations included the number of effective migrants, the number of effective parents, total number of individuals, the sex ratio, and the number of adults. The number of migrants in the population and the proportion of migrant SNPs ($nSNP_M$) allowed us to monitor how migrant alleles were distributed across individuals. Genetic diversity and fitness were classified as the observed heterozygosity, calculated across all SNPs ($nSNP_D$ + $nSNP_M$) and in drift SNPs ($nSNP_D$). Inbreeding in the population ($F_{IS}$) and the amount of divergence in the recipient population ($F_{ST}$) at that year as compared to the source population and compared to the initialized recipient population were evaluated using the *hierfstat* package in R. After all simulation years, all dead individuals were written back into the population to calculate the lifetime (LRS) of all individuals by evaluating the number of mates, number of offspring, and the number of offspring that survived to maturity for all individuals.

Quantitative Comparisons

We compared the effect of all parameters on genetic diversity and population persistence by comparing the output of the simulated populations to control simulations. For each parameter set, we measured the heterozygosity and the distribution of each SNP type to examine how migration altered the genetic diversity of the recipient population. Additionally, we evaluated pairwise $F_{ST}$

values each year compared to the initialized recipient and source populations. Population size, inbreeding level ($F_{IS}$), and lifetime reproductive success were also calculated. We ran 100 replicates for each combination of parameters, with each simulation run in R on a high-performance computing cluster.

**Supplementary Figure 1.** Genetic and demographic responses to migration when migrants are (orange) and are not (grey) preferentially chosen as mates with a population crash to 70% of the historical population size. Movement of a single individual per year (solid line), burst migration of 100 individuals once (year 151; dashed line), and four pulse migrations of 25 individuals (years 151, 165, 181, 195; dashed and dotted line) was used to support and bolster these populations through remediation (year 151) and population recovery. These trends were compared to the same demographic patterns but without migration (dotted line). The proportion of migrant ancestry present in the recipient populations (A), observed heterozygosity (B), divergence of the recipient populations from the historical populations over time (C), and divergence of the recipient populations from the migrant source populations each year (D) illustrate the new evolutionary trends resulting from these migration decisions. Lines represent mean values across 100 replicates and error bars represent the confidence intervals needed to compare among parameter sets assuming alpha = 0.05 (i.e., 95% confidence intervals).

**Supplementary Figure 2.** Proportion of populations that remained viable and survived to the completion of the simulation when the population was critically endangered among various migration rates. Vertical grey line indicates the year of supplementation for burst migrations and the first pulse immigration. All other simulations resulted in 100% viability.

**Supplementary Figure 3.** Inbreeding level ($F_{IS}$) in the absence of migration (A), with one migrant per generation (B), and with burst (C) and pulse (D) migrations compared among extinction risk categories (critically endangered, 90% reduction; endangered, 70% reduction; vulnerable, 30% reduction; no population reduction). Note that the y-axis differs within this figure. Grey vertical lines depict the years at the start of population decline (y = 100) and subsequent habitat restoration (y = 150). The black horizontal line shows when $F_{IS}$ is zero. Lines represent mean values across 100 replicates and shaded areas represent the confidence intervals needed to compare among parameter sets assuming alpha = 0.05 (i.e., 95% confidence intervals).

**Supplementary Figure 4.** Sex ratio (females : males) in the absence of migration (A), with one migrant per generation (B), and with burst (C) and pulse (D) migrations compared among extinction risk categories (critically endangered, 90% reduction; endangered, 70% reduction; vulnerable, 30% reduction; no population reduction). Grey vertical lines depict the years at the start of population decline (y = 100) and subsequent habitat restoration (y = 150). The black horizontal line depicts a 50:50 sex ratio; the population is female dominated when the ratio < 0.5 and male dominated when the ratio > 0.5. Lines represent mean values across 100 replicates and shaded areas represent the confidence intervals needed to compare among parameter sets assuming alpha = 0.05 (i.e., 95% confidence intervals).

**Supplementary Figure 5.** Lifetime reproductive success in the absence of migration (A), with one migrant per generation (B), and with burst (C) and pulse (D) migrations compared among extinction risk categories (critically endangered, 90% reduction; endangered, 70% reduction; vulnerable, 30% reduction; no population reduction). Lines represent mean values across 100 replicates and shaded areas represent the confidence intervals needed to compare among parameter sets assuming alpha = 0.05 (i.e., 95% confidence intervals).
