## Supplementary figures and images for "Habitat remediation followed by managed connectivity reduces unwanted changes in evolutionary trajectory of high extirpation risk populations"

### Supplemental Figure 1

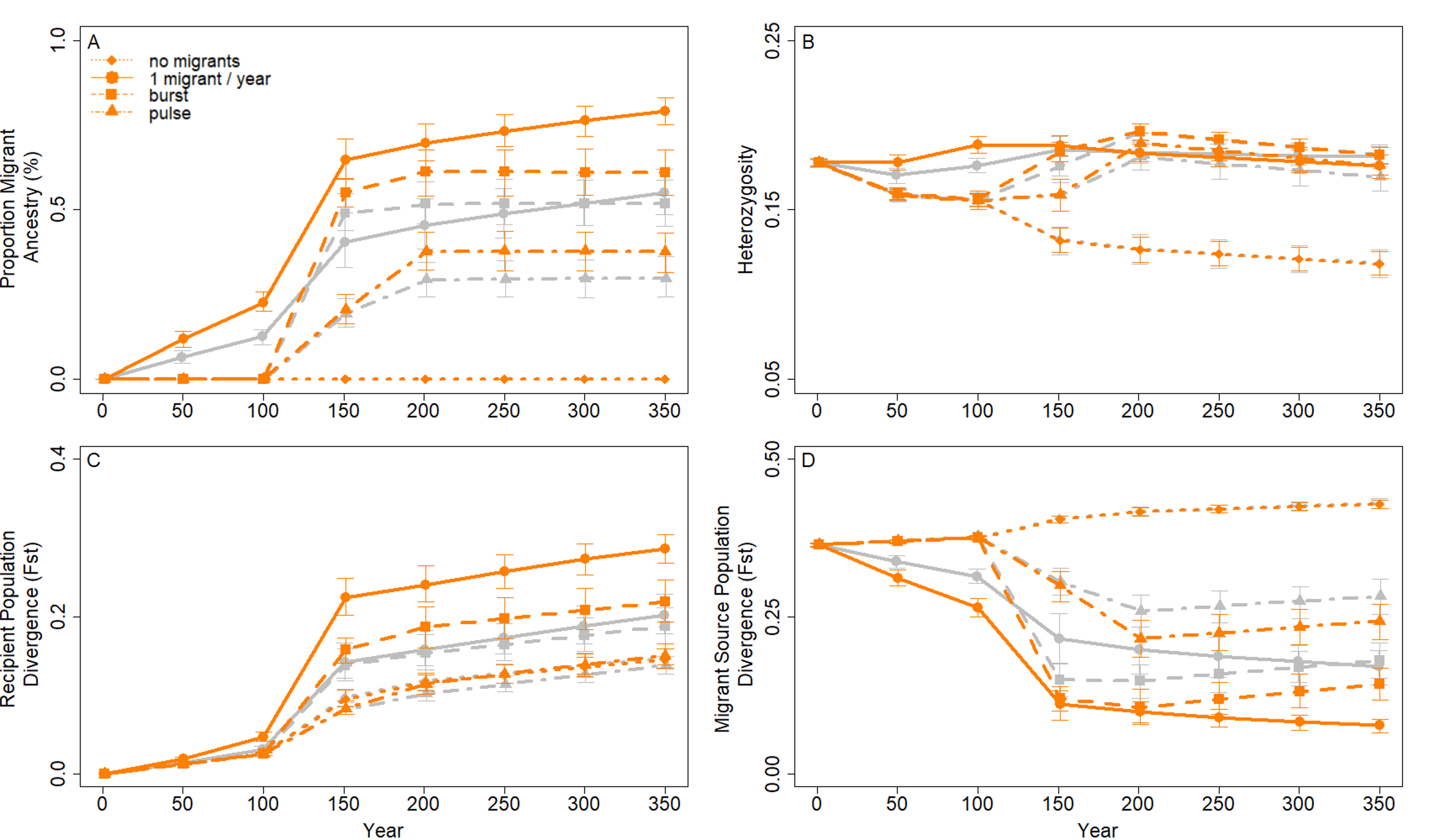

### Supplemental Figure 2

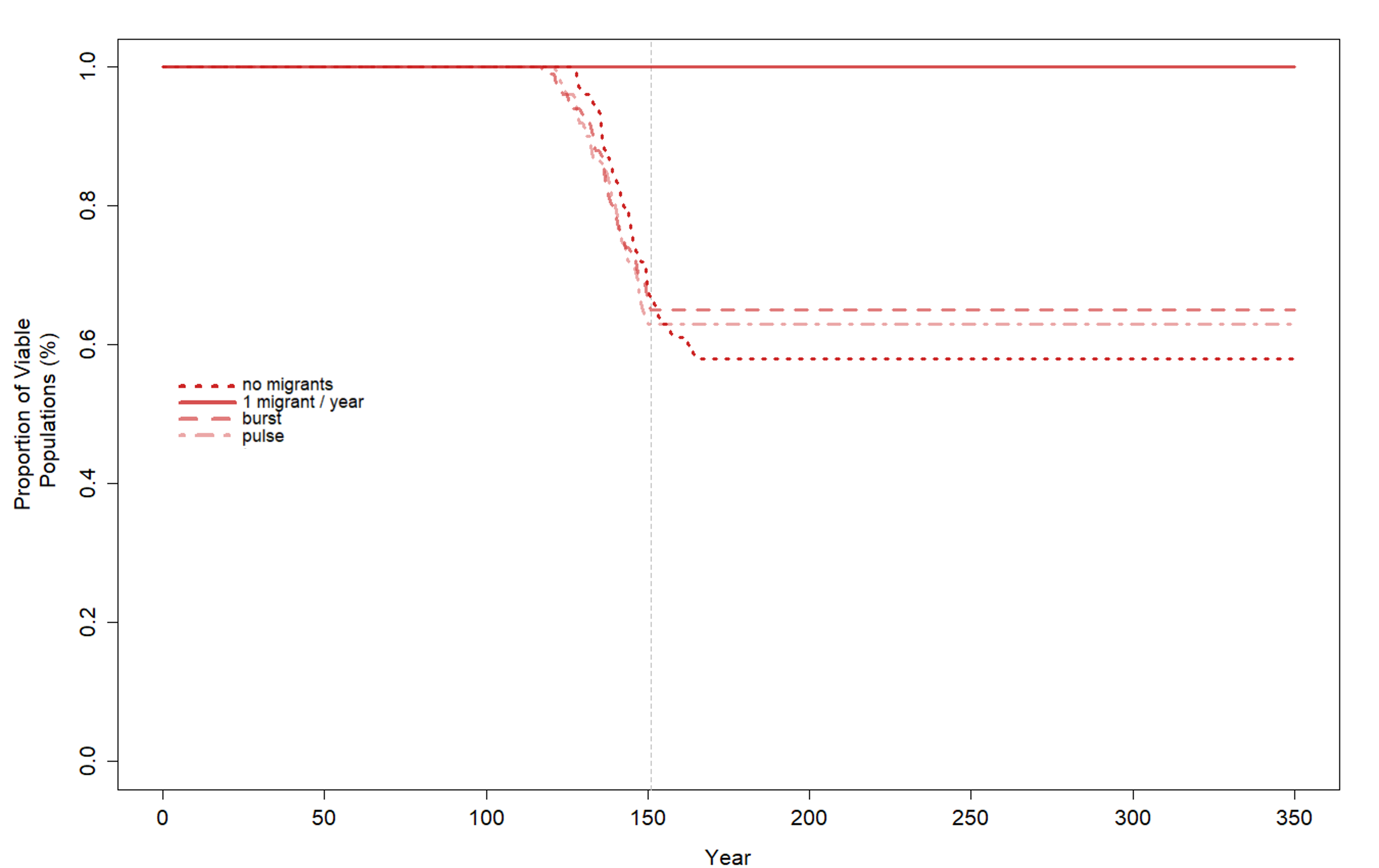

### Supplemental Figure 3

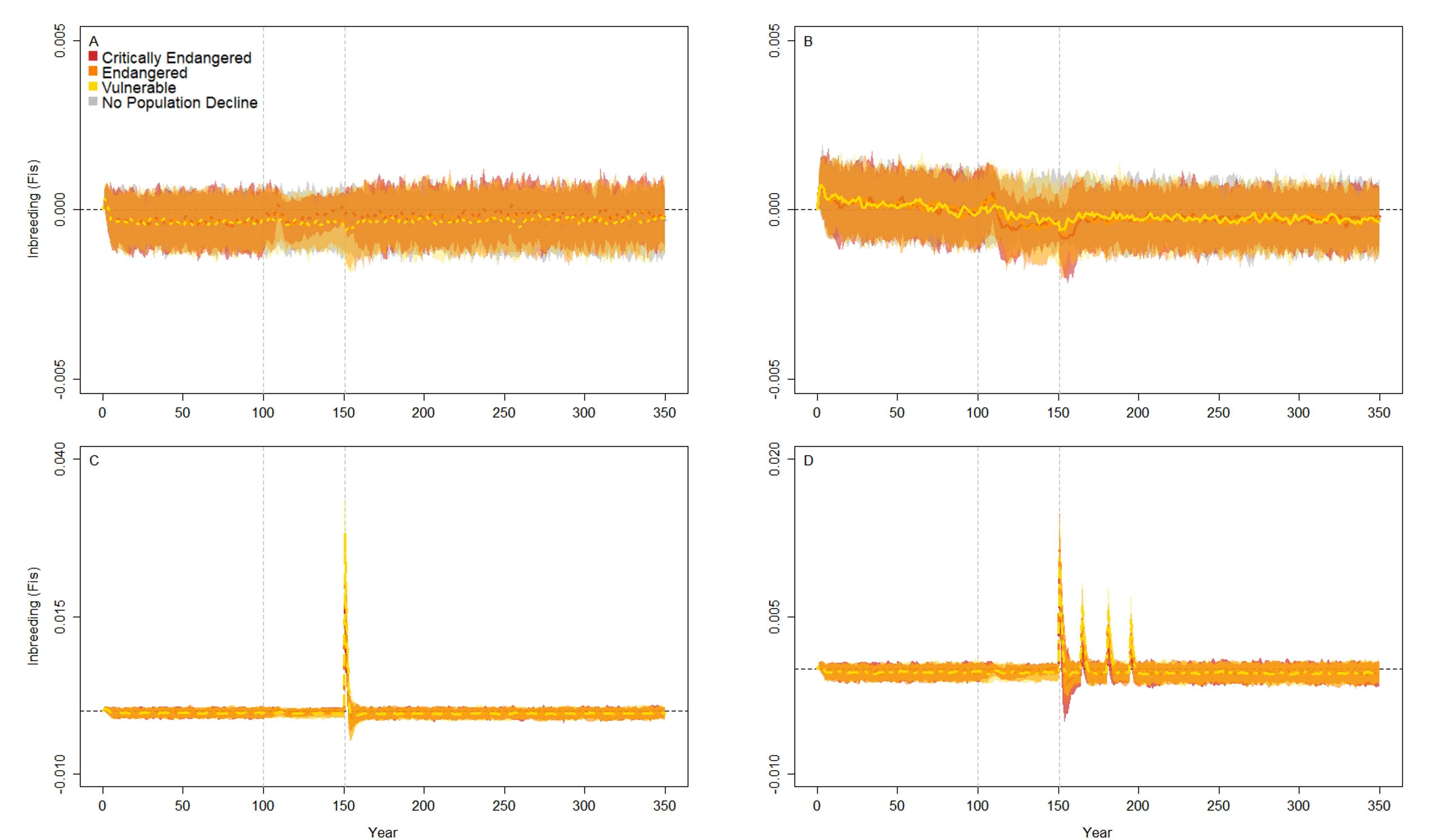

### Supplemental Figure 4

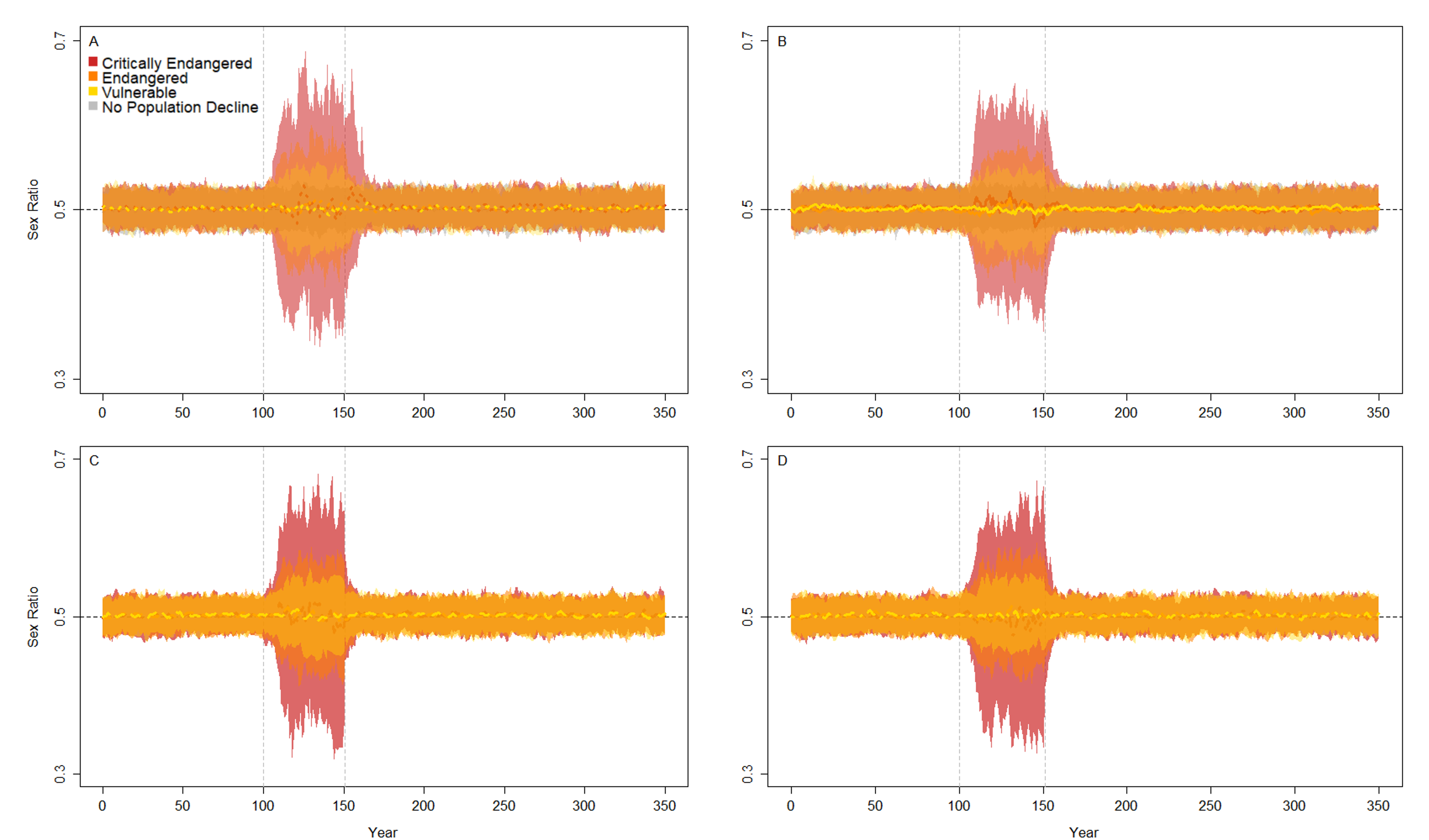

### Supplemental Figure 5

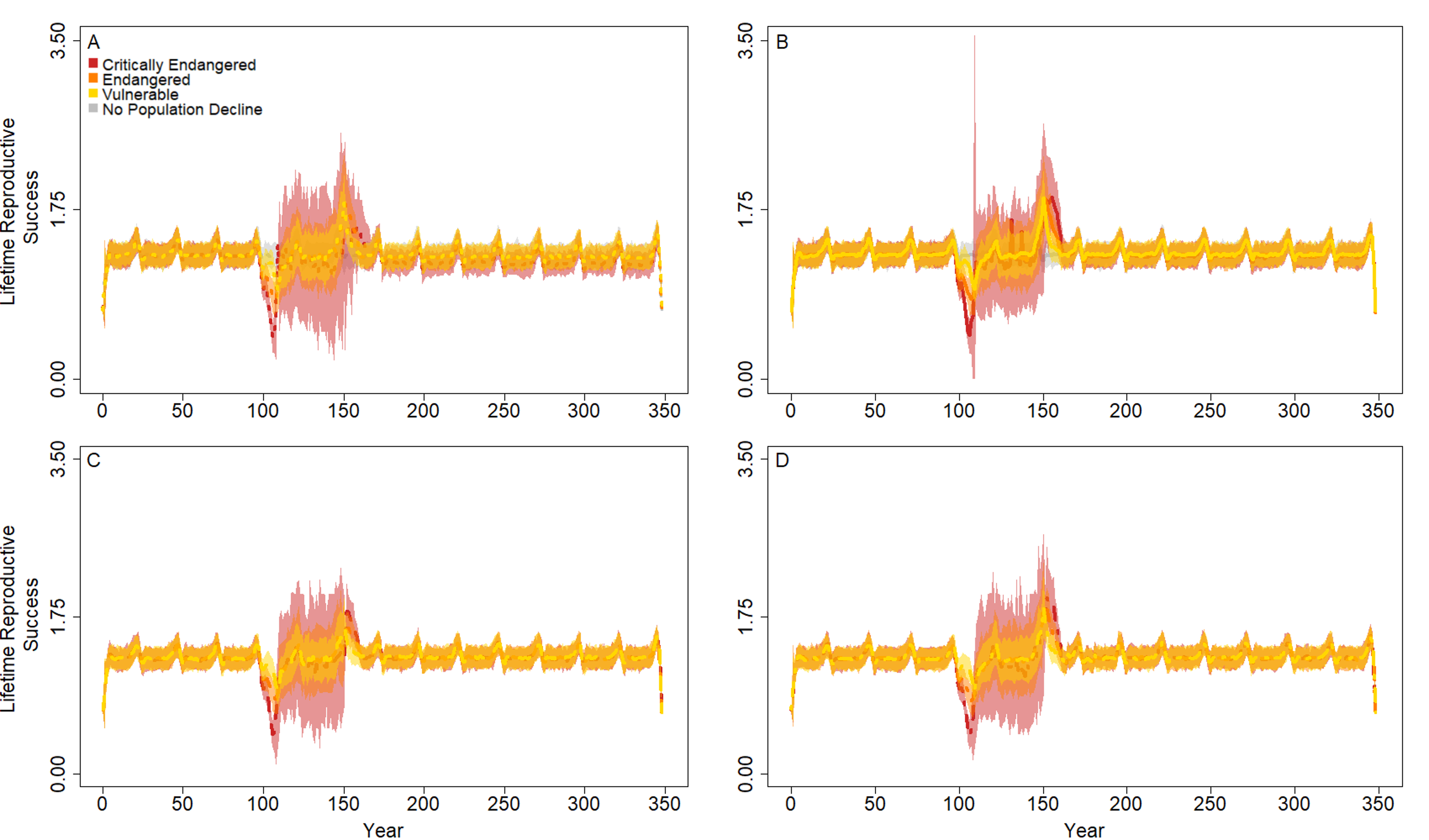
